# Microglia are necessary for toxin-mediated demyelination and activation of microglia is sufficient to induce demyelination

**DOI:** 10.1101/501148

**Authors:** Dave E. Marzan, Brian L. West, James L. Salzer

## Abstract

Microgliosis is a prominent pathological feature in many neurological diseases including multiple sclerosis (MS). The precise role of microglia during demyelination, and the relative contributions of microglia vs. peripheral macrophages, are incompletely understood. Here, using a genetic fate mapping strategy, we identify microglia as predominant responders and key effectors of demyelination in the cuprizone (CUP) model. Pharmacological depletion of microglia demonstrates these cells are necessary for the demyelination, loss of oligodendrocytes, and reactive astrocytosis normally evident in this model. Electron microscopy (EM) and serial block face imaging show myelin sheaths remain intact in CUP treated mice depleted of microglia. However, these damaged myelin sheaths are lost upon-repopulation of microglia. Injection of colony-stimulating factor-1 to drive focal microgliosis in white matter is sufficient to induce focal demyelination *in vivo*. These studies indicate activated microglia are required for demyelination that results from primary myelin pathology and are sufficient to induce demyelination directly.

## INTRODUCTION

Microglia are the resident tissue macrophages of the central nervous system (CNS) comprising ∼10% of cells in the adult mouse and human brain (Lawson LJ, 1990; Mittelbronn, Dietz, Schluesener, & Meyermann, 2001). These highly dynamic cells derive from yolk sac precursors that colonize the brain early during embryonic development and are maintained autonomously throughout adult life by clonal expansion (Ajami, Bennett, Krieger, Tetzlaff, & Rossi, 2007; Davalos et al., 2005; Ginhoux et al., 2010; Tay et al., 2017). In the healthy brain, microglia are constantly changing their morphology to surveil the brain parenchyma, remodel synapses during development, contribute to motor learning and memory, and phagocytose dead cells (Fourgeaud et al., 2016; Nimmerjahn, Kirchhoff, & Helmchen, 2005; Parkhurst et al., 2013; Schafer et al., 2012; Sierra et al., 2010). Microglia are also increasingly appreciated to have central roles in a wide variety of neurological disorders including Alzheimer’s disease, neuropathic pain, and adult-onset leukodystrophy with axonal spheroids and pigmented glia (ALSP) (Beggs, Trang, & Salter, 2012; Chitu et al., 2015; Derecki et al., 2012; Frautschy et al., 1998; Tsuda et al., 2003).

The role of microglia in myelin disorders is of intense interest, in particular in multiple sclerosis (MS), a progressive autoimmune inflammatory and demyelinating disease and the most common cause of neurological disability in young adults (Compston A, 2002). While the etiology of MS still remains to be established, pathologically it is characterized by focal inflammatory lesions that result in demyelination, axonal loss and gliosis (Lassmann, 2018). Classically, the inflammation is thought to result from peripheral immune cell infiltration into central nervous system (CNS) (Fletcher, Lalor, Sweeney, Tubridy, & Mills, 2010). As tools for identifying and imaging microglia have advanced, numerous studies have found reactive microglia in both active, and pre-active lesions suggesting microglia contribute to both the development and progression of MS (Lassmann, 2014; van der Valk & Amor, 2009; Z. Zhang et al., 2011).

The roles of microglia in demyelination have been studied in several murine models of demyelination. These models include cuprizone (CUP), a chemical toxin, which is provided in the diet of adult mice. CUP treatment results in reliable and preferential demyelination of the corpus callosum (CC) that is followed by robust remyelination that ensues upon cessation of cuprizone treatment (Matsushima G, 2001; Praet, Guglielmetti, Berneman, Van der Linden, & Ponsaerts, 2014; Wergeland, Torkildsen, Myhr, Mørk, & Bø, 2012). Using this and other models of demyelination, several studies have investigated the role of microglia in demyelination. Some earlier studies reported that attenuating microglia activity reduces demyelination (McMahon, Cook, Suzuki, & Matsushima, 2001; Skripuletz et al., 2010; Wergeland et al., 2011) whereas others found microglia derived signals reduce demyelination and promote remyelination (Heather A. Arnett et al., 2002; H. A. Arnett et al., 2001; Olah et al., 2012). Difficulties in distinguishing microglia from infiltrating macrophages using standard markers (e.g. Iba1, CD11b) may contribute to these earlier conflicting reports as microglia and infiltrating macrophages share expression of many cell surface markers; their marker expression and morphologies are also sensitive to experimental manipulation (Bogie, Stinissen, & Hendriks, 2014; Elkabes, DiCicco-Bloom, & Black, 1996; Graeber et al., 1998). In addition, when macrophages enter the CNS, they can acquire expression of markers considered to be microglia-specific further confounding analysis (F. C. Bennett et al., 2018).

Here, we have used a genetic strategy to differential label microglia and macrophages as a first step in assessing their roles during demyelination in the CUP model. We show CNS resident microglia markedly and selectively expand during demyelination induced by CUP. In complementary studies in which we pharmacologically depleted microglia with a colony stimulating factor 1 receptor (CSF1R) inhibitor, PLX3397 (PLX) (Tap et al., 2015), we further demonstrate that small numbers of surviving microglia are capable of rapidly repopulating the CNS including that driven by demyelination. By pre-depleting microglia with PLX and then inducing demyelination, we demonstrate demyelination is markedly reduced indicating that microglia are required for the demyelination induced by CUP. Finally, by inducing focal microgliosis in the corpus callosum, we provide direct evidence that activation of microglia is sufficient to induce demyelination. These results provide compelling evidence for the key role of microglia in mediating demyelination in CUP and by analogy, in other demyelinating diseases.

## RESULTS

### Fate mapping identifies microglia, not macrophages, at sites of CUP-mediated demyelination

An increase in the numbers of microglia/macrophages during demyelination in the CUP model has long been known (Blakemore, 1973; Matsushima G, 2001). However, the standard antibody markers used to stain microglia (e.g. Iba1, CD11b) do not distinguish between microglia and peripheral macrophages, leading to uncertainty with regard to their origins and designation (Hiremath et al., 1998). To obviate limitations of traditional immunofluorescence, we utilized a fate mapping strategy to differentially label microglia vs. macrophages *in vivo* by using the *CX3CR1*^*CreER-iresGFP*^;*Rosa26*^*stop-DsRed*^ mouse line. In this system, CNS microglia retain their DsRed expression 30 days after tamoxifen treatment whereas peripheral macrophages do not and only express GFP under control of the CX3CR1 promoter (Parkhurst, et al., 2013), allowing confident assessment of their contributions in the inflamed brain.

We used this differential labeling strategy to identify the source of CD11b^+^ cells found in the CC during CUP mediated demyelination (Supplemental Fig. 1). Four-week old *CX3CR1*^*CreER-iresGFP/+;*^*Rosa26*^*stop-DsRed*^ mice were treated with tamoxifen and 30 days later were placed on a CUP diet (Fig. 1A). Mice were sacrificed at 3 and 5 weeks of CUP (Fig. 1B), timing associated with early and peak of demyelination, respectively. Close examination of the CC of *CX3CR1*^*CreER-iresGFP/+*^*;Rosa26*^*stop-DsRed*^ mice on control and CUP (3 and 5 week) diets revealed the great majority of all labeled cells on control and CUP diets (>99%) were DsRed^+^ microglia and virtually all of these were also GFP^+^ in mice (Fig. 1B and C). DsRed^+^/GFP^-^ cells made up less than ∼1% of all cells at all time points; these presumably reflect microglia with weak/undetectable GFP expression. Of note, macrophages i.e. DsRed^-^/GFP^+^ cells, were rare and comprised less than 0.5% of all cells in the CC of control and CUP treated mice (Fig. 1C).

**Figure 1.**
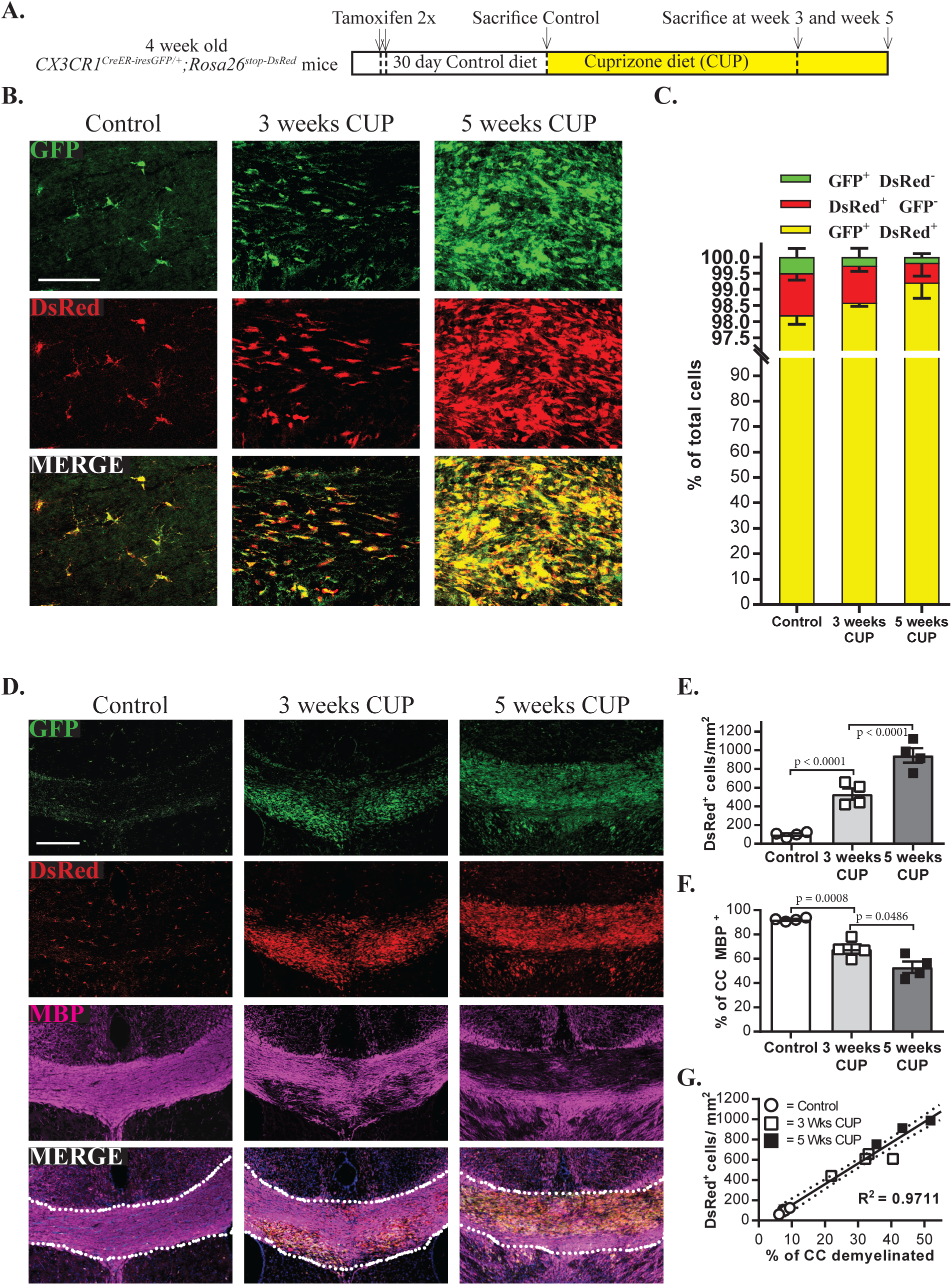
Genetic labeling identifies microglia in the cuprizone model of demyelination. A. Schematic of the experimental design. *CX3CR1*^*CreER-iresGFP/+*^*;Rosa26*^*stop-DsRed*^ mice were treated with tamoxifen, and then maintained for an additional month to ensure turnover of macrophages. Microglia remain DsRed-positive; macrophages are DsRed-negative at this time. Mice were then placed on a cuprizone diet (CUP) for 3 or 5 weeks before analysis.
B. Confocal micrographs of coronal brain sections of CUP treated mice stained for GFP and DsRed demonstrates virtually all cells are GFP and DsRed positive microglia. Scale bar = 100 μm.
C. Quantification of fate mapped cells following cuprizone treatment. Bar graphs showing the distribution of GFP^+^ DsRed^−^ cells (green), GFP^-^ DsRed^+^ cells (red), and GFP^+^ DsRed^+^ double positive cells (yellow) indicate nearly all cells are microglia. N= 4 per group; data are mean +/- s.e.m.; Student’s t-test.
D. Confocal micrographs of coronal brain sections stained for GFP, DsRed, and MBP show microgliosis and demyelination are tightly correlated. Scale bar = 200 μm.
E. Quantification of the numbers of microglia (i.e. DsRed^+^ cells) in the CC of mice on control diet, 3 and 5 weeks of CUP. Each symbol represents 1 animal. N=4 per group; data are mean +/- s.e.m.
F. Quantification of the percentage of CC positive for MBP for mice on control diet and 3 and 5 weeks of CUP. Each symbol represents 1 animal. N=4 per group; data are mean +/- s.e.m.
G. Linear regression correlating microglia to demyelination in the CC based on quantifications in Figs. 1E and F; R^2^ = 0.9711.

These data indicate that microglia, not infiltrating macrophages, are the source of monocytes in demyelinated lesions of CUP treated mice and raise the possibility that they contribute to demyelination in this model. In potential agreement, we found a close correlation between microglia numbers and the extent of demyelination. Thus, areas of peak demyelination directly corresponded to sites of high microglia density as seen in representative confocal micrographs (Fig. 1D). The numbers of microglia increased from 98.0 ± 13.4 DsRed^+^ cells/mm^2^ in the CC of mice on the control diet to 531.7 ± 59.7 cells/mm^2^ at three weeks of CUP and to 944.1 ± 76.8 cells/mm^2^ at five weeks of CUP (Fig. 1E). Levels of myelin in corresponding sections of the CC, as determined by staining for MBP, decreased in proportion to the increased numbers of microglia, from 92.3 ± 0.7% in controls, to 68.1 ± 3.8% and 53.1 ± 4.7% at three and five weeks of CUP, respectively (Fig. 1F). Linear regression shows a strong statistical correlation (R^2^ = 0.9711) between the numbers of DsRed^+^ cells/microglia and the extent of demyelination (Fig. 1F). A similar correlation between increased numbers of microglia and demyelination in WT C57BL/6J mice on control and cuprizone diets was also observed based on CD11b and MBP immunoreactivity (Supplemental Fig. 1E).

### Depletion of microglia delays demyelination

To examine the role of microglia in the CUP model of demyelination, we utilized a well-established, pharmacological strategy of depleting microglia resulting from treatment with the CSF1R inhibitor PLX3397 (PLX) (M. R. P. Elmore, Lee, West, & Green, 2015; Monica R. P. Elmore et al., 2014). Treatment with PLX is a highly effective and well tolerated strategy, with near complete elimination of microglia in the healthy brain after ∼ 1 week. Cessation of PLX treatment is followed by a rapid (∼ 1 week) rebound in microglia numbers. By fate-mapping in *CX3CR1*^*CreER-iresGFP/+*^*;Rosa26stop*^*-DsRed*^ mice, we confirmed that these repopulating microglia arise from the small numbers of surviving microglia that persist during PLX treatment (Supplemental Fig. 2), in agreement with a recent report (Huang et al., 2018).

We next examined the role of microglia in the CUP model of demyelination by combining CUP (0.2%) and PLX3397 (290 mg/kg) into a single chow, i.e. CUP/PLX. *CX3CR1*^*CreER-iresGFP/+;*^*Rosa26*^*stop-DsRed*^ mice were fed diets containing CUP, PLX, or CUP/PLX (Fig. 2A). Brains of mice on control diets or 3 and 5 weeks of CUP, were stained for DsRed, GFP, and MBP (Fig. 2B). Mice on 3 weeks of CUP/PLX diet had fewer DsRed^+^ cells (254.4 ± 39.3 cells/mm^2^) compared to mice fed CUP for 3 weeks (514.1 ± 35.3 cells/mm^2^; p=0.0050) (Fig. 2C). Of particular interest, PLX treatment resulted in less demyelination at this early timepoint. Thus, mice fed CUP alone for 3 weeks had 56.9 ± 5.7% of CC positive for MBP compared to controls whereas mice on CUP/PLX diet were much higher, 94.9 ± 2.2% than controls (p=0.0066; Fig. 2E). Taken together, these results strongly implicate microglia in demyelination in the CUP model.

**Figure 2.**
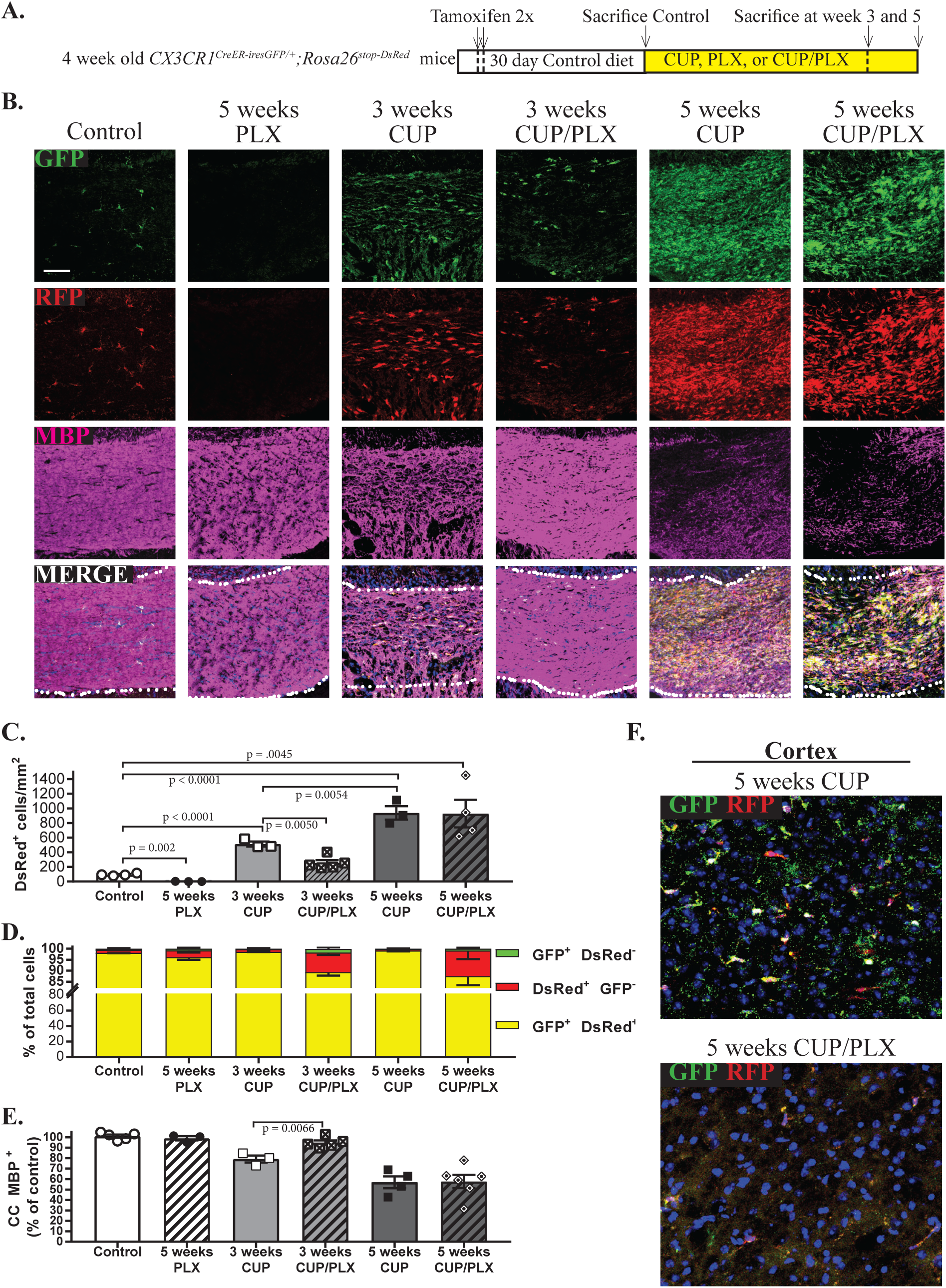
Pharmacological depletion of microglia delays demyelination. A. Schematic of the experimental design. *CX3CR1*^*CreER-iresGFP/+*^*;Rosa26*^*stop-DsRed*^ mice were treated with tamoxifen, and after an additional month were placed on PLX for 5 weeks, cuprizone (CUP) for 3 or 5 weeks, or CUP/PLX for 3 or 5 weeks before analysis.
B. Confocal micrographs of coronal brain sections of cuprizone treated mice stained for GFP and DsRed demonstrates virtually all cells are double positive microglia. Scale bar = 100 μm. Microglia are nearly absent after PLX treatment. Microglia increase with CUP treatment but not at 3 weeks of CUP/PLX combination diet. Demyelination is closely correlated to the levels of microglia. Scale bar = 100 μm.
C. Total numbers of DsRed^+^ microglia/mm^2^ in different treatment groups are shown. Groups include mice on control diet (N=4), 5 weeks PLX (N=3), 3 weeks CUP (N=3), 3 weeks CUP/PLX (N=5), 5 weeks CUP (N=3) and 5 weeks of CUP/PLX (N=4). Each symbol represents 1 animal; data are mean +/- s.e.m.
D. Quantification of the proportion of microglia, i.e. DsRed^+^ (red) and GFP^+^ DsRed+ double positive (yellow) cells vs. macrophages, i.e. GFP^+^ DsRed-(green) cells; groups are the same as those shown in panel C.
E. Quantification of MBP levels in the CC, graphed relative to control diet mice. Each symbol represents 1 animal; data are mean +/- s.e.m.
F. Representative micrograph from the cortex of mice shown in panel B on 5 weeks of CUP or on 5 weeks of CUP/PLX diet stained for GFP, DsRed, and Hoechst (blue). While large numbers of microglia are present in the CC of mice on CUP/PLX at 5 weeks, they are largely depleted in the cortex.

PLX treatment was less effective at depleting microglia and reducing demyelination at later time points. At 5 weeks as mice on CUP and CUP/PLX had comparable numbers of DsRed^+^ cells (939.5 ± 92.2 vs 929.8 ± 188.1 cells/mm^2^, respectively) and the extent of demyelination was similar (56.91 ± 5.70% vs 57.62 ± 6.28 % of CC MBP^+^, respectively) (Figure 2C and E). Population analysis of fate mapped cells in mice fed CUP/PLX diet confirmed the great majority of labeled cells were *bona fide* microglia; only 1.7 ± 0.6% and 0.7 ± 0.4% of cells were macrophages, i.e. DsRed^-^/GFP^+^ after 3 and 5 weeks of treatment, respectively Thus, the cells that repopulated the CC arose from residual microglia. While microglia persisted/expanded with demyelination of the CC on CUP/PLX, they were effectively eliminated from all other sites in the brain including the cortex at these later time points (Fig. 2F). These results demonstrate that PLX is effective at decreasing microglia numbers and demyelination early, yet loses its efficacy to deplete CC microglia and attenuate demyelination at late stages of the CUP model.

To further corroborate our findings of microglial persistence and repopulation in mice at 5 weeks of the CUP/PLX diet, we utilized mice deficient for the C-C chemokine receptor type 2 (CCR2) (Supplemental Fig. 3). CCR2 is required for monocyte/macrophage recruitment to the inflamed CNS (Ajami, Bennett, Krieger, McNagny, & Rossi, 2011; Boring et al., 1997). We treated CCR2^-/-^ mice with CUP and CUP/PLX diets and assessed CD11b cell numbers and the levels of demyelination (Supplemental Fig. 3). There was no difference in CD11b^+^ cells in the CC of WT vs CCR2^-/-^ mice on CUP and CUP/PLX diets at 3 and 5 weeks (Supplemental Fig. 3C). Furthermore, myelin levels were comparably reduced in the CC of CCR2^-/-^ mice and WT mice on CUP/PLX at both 3 and 5 weeks (Supplemental Fig. 3D). These findings corroborate the fate mapping in the *CX3CR1*^*CreER-iresGFP/+;*^*Rosa26*^*stop-DsRed*^ mice that indicate macrophages do not infiltrate or contribute to demyelination in the CUP model.

### Prophylactic depletion of microglia blocks CUP-mediated demyelination

To limit the rapid repopulation of microglia in the CC of CUP/PLX mice and characterize further the role of microgliosis in CUP-mediated demyelination, we pretreated *CX3CR1*^*CreER-iresGFP/+;*^ *Rosa26*^*stop-DsRed*^ mice with PLX for two weeks prior to starting them on CUP/PLX for 5 weeks. These were compared to the same mice treated for 5 weeks with CUP alone (Fig. 3A). We also examined mice on seven weeks of the PLX diet alone, which served as a control for chronic depletion of microglia and/or possible off target effects of the CSF1R inhibitor. Sections of the CC were stained for GFP, DsRed and MBP (Fig. 3B). Microgliosis, as determined by DsRed^+^ labeled cells, was substantially reduced by the two-week pretreatment with PLX followed by 5 weeks of PLX/CUP, i.e. 124.9 ± 11.6 cells/mm^2^ vs. 917.1 ± 75.3 cells/mm^2^, respectively (Fig. 3C). Of note, the mice pre-treated with PLX also exhibited much less demyelination in the CC compared to mice fed CUP alone, i.e. 76.8 ± 4.1% vs. 52.8 ± 7.2%, respectively (Fig. 3D).

**Figure 3.**
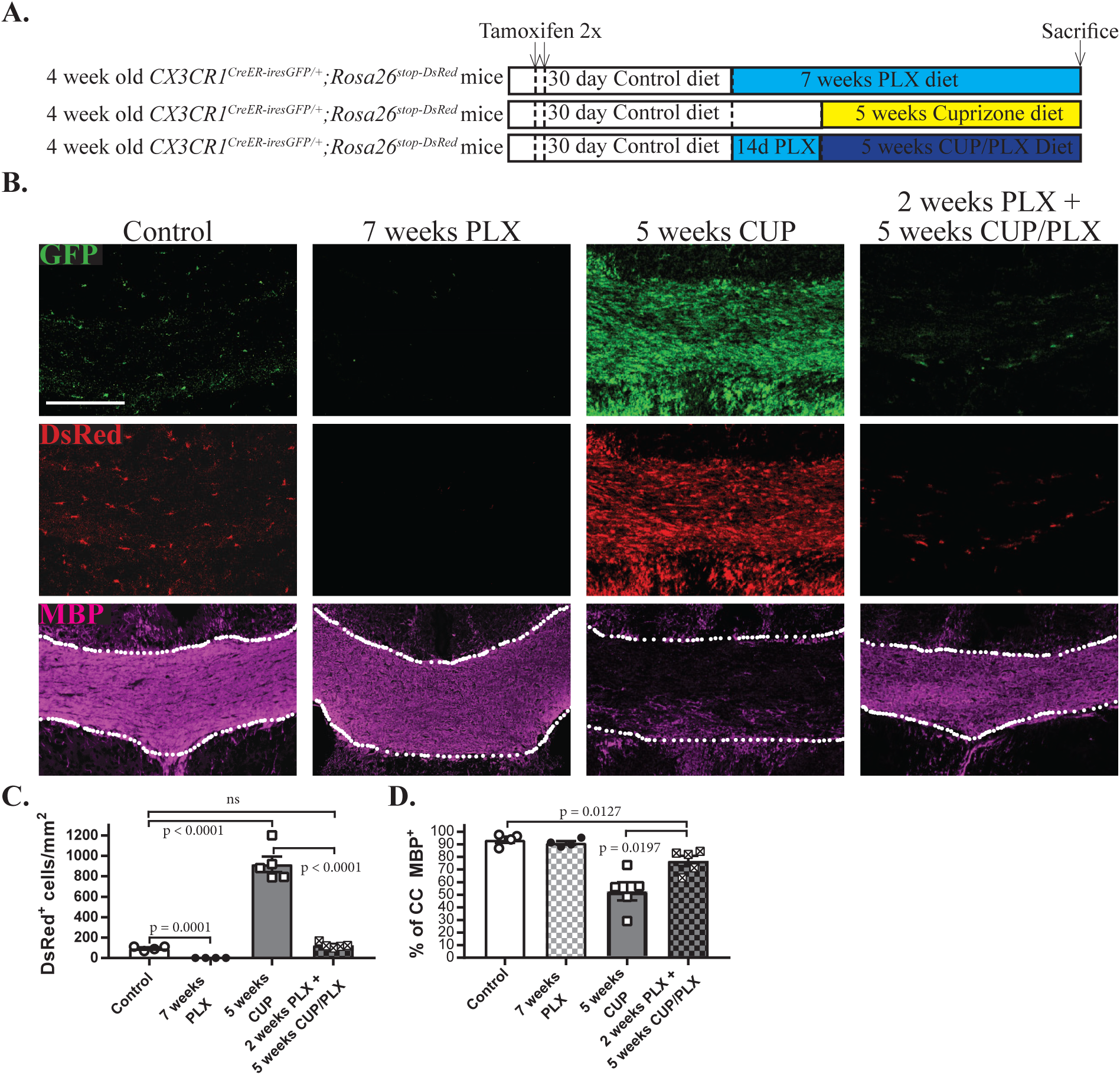
Prophylactic depletion of microglia blocks CUP-mediated demyelination. A. Schematic of the experimental design. *CX3CR1*^*CreER-iresGFP/+*^*;Rosa26*^*stop-DsRed*^ mice were treated with tamoxifen. After an additional 6 weeks, one group of mice was placed on CUP for 5 weeks. Other mice were treated with tamoxifen and after an additional month received either PLX for 7 weeks or PLX for 2 weeks followed by CUP/PLX for 5 weeks.
B. Confocal micrographs from the CC of the experimental groups indicated were stained for GFP, DsRed, and MBP. Pretreatment with PLX followed by ongoing PLX abrogates CUP mediate microgliosis and substantially attenuates demyelination. Scale bar = 200 μm.
C. Quantification of DsRed^+^ microglia/mm^2^ in the CC of mice on control diet (N = 4), 7 weeks PLX (N = 4), 5 weeks of CUP (N = 5) and 2 weeks PLX + 5 weeks CUP/PLX (N = 4). Each symbol represents 1 animal; data are mean +/- s.e.m.
D. Quantification of MBP levels in the CC of the mouse groups shown in Fig. 3C. Each symbol represents 1 animal; data are mean +/- s.e.m.

These results strongly suggest that in the absence of microglia, myelin is preserved despite cuprizone treatment. However, these results do not exclude the possibility that MBP staining detects myelin debris that is not cleared when microglia are depleted, rather than myelin sheaths. We therefore examined the integrity of the myelin sheaths directly via electron microscopy of the CC of mice treated with cuprizone for 5 weeks, treated with the combination diet (i.e. CUP/PLX) for 5 weeks, or treated with 2 weeks of PLX then 5 weeks of the combination diet (Fig. 4). In general, mice on the CUP diet exhibited large areas of demyelination, with the majority of axons lacking ensheathment; there were also modest numbers of microglia and astrocytes present (Fig. 4A). In mice on the CUP/PLX diet, many fields appeared to be actively demyelinating with evidence of fragmented myelin sheaths; this is consistent with the rebound of microglia between weeks 3 and 5 (Fig. 2). Microglia were frequently present, typically laden with intracellular vesicles containing partially degraded myelin, with lamellae evident at high power (Fig. 4G). In contrast, axons in mice pre-treated with PLX appeared myelinated similarly to controls (Fig. 4A). Quantification of the numbers of myelinated and unmyelinated axons (Fig. 4D) corroborates preservation of myelin sheaths in the absence of microglia despite cuprizone treatment.

**Figure 4.**
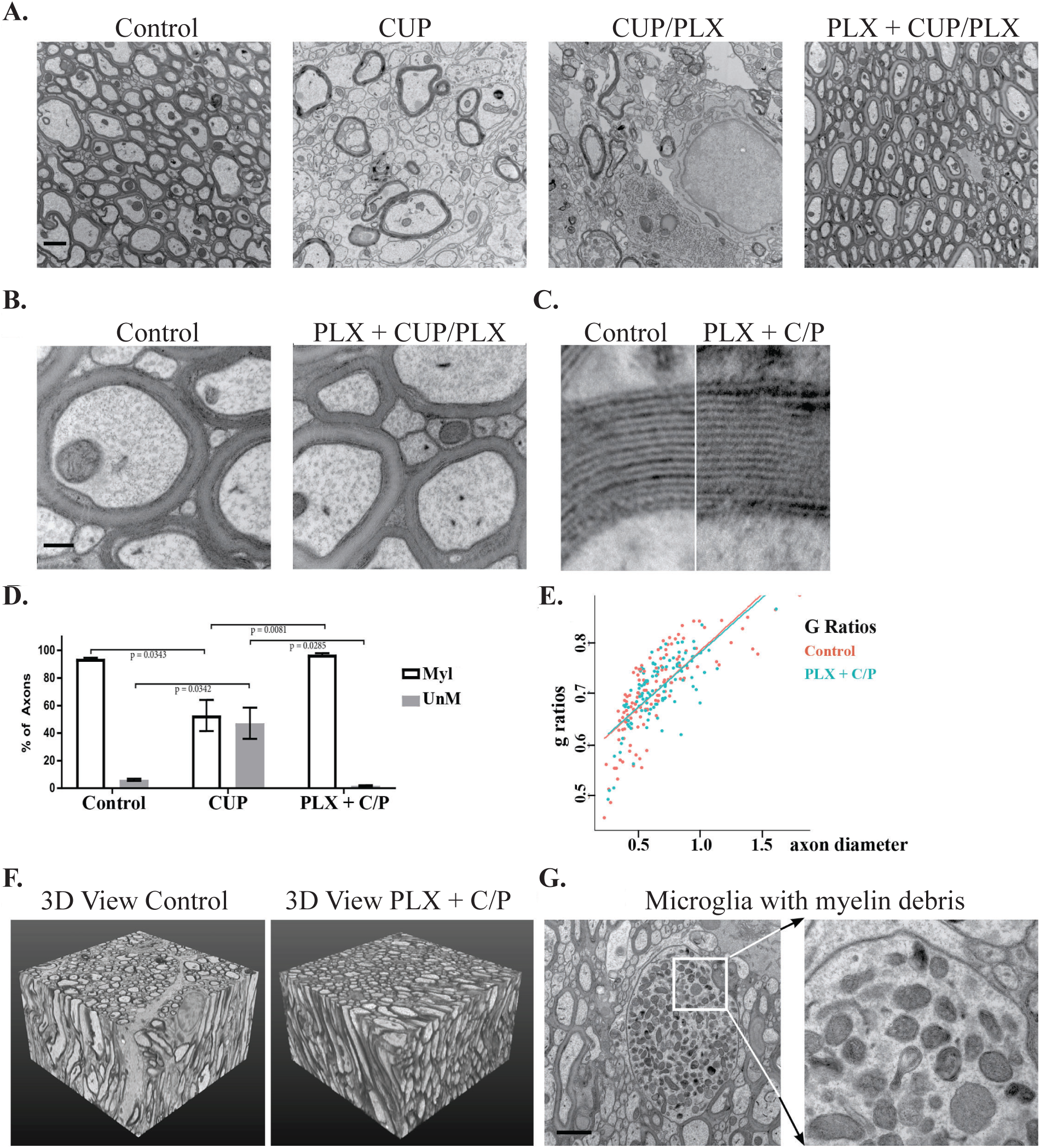
Electron microscopic analysis of cuprizone-treated myelin in the presence or absence of microglia. A. Electron micrographs of the CC of mice on control diet, CUP (5 weeks), CUP/PLX (5 weeks), and PLX for 2 weeks then PLX/CUP (5 weeks). There is substantial demyelination of the CC of mice on CUP and CUP/PLX vs. near complete protection of myelin in the PLX + CUP/PLX cohort. Scale bar = 2 μm.
B. Comparison of the morphology of myelin sheaths in controls vs. the PLX + CUP/PLX cohort. The persistent myelin in the latter group appears nearly normal. Scale bar = 0.2 μm
C. Comparison of the morphology of myelin sheaths in controls vs. those that persist in the PLX + CUP/PLX cohort show myelin spacing is similar, although the major dense lines are less pronounced.
D. Quantification of the percentage of myelinated vs. unmyelinated axons in the CC of mice on control diet, treated with cuprizone for 5 weeks, or treated with PLX for 2 weeks then CUP/PLX for 5 weeks.
E. G ratios of myelin sheaths in controls vs. those that persist in the PLX + CUP/PLX cohort.
F. 3D reconstructions of serial block face EM sections taken from the body of the CC of control and PLX + CUP/PLX treated mice. Volume show is 16 × 16 × 10 μm. Animated reconstructions of serial block face EM sections are shown in the supplemental data.
G. An example of a microglial cell filled with vesicles in a partially demyelinated field of the CC of mice treated with CUP/PLX for 5 weeks. Scale bar = 1 μm. The higher power image (boxed) shows that many of these vesicles contain partially degraded myelin, whose lamellae are still visible.

We compared myelin in controls to that in mice pre-treated with PLX then CUP/PLX in further detail. The preserved, cuprizone treated myelin appeared subtly different, with less oligodendrocyte cytoplasm in the inner turn; sheaths appeared more tightly packed with less extracellular space. In addition, the characteristic alternating dark (major dense) and light (intraperiod) lines were less distinct (Fig. 4C). Myelin lamellar spacing (Fig. 4C) and g ratios (Fig. 4E) were comparable. Serial block face reconstructions and animations revealed remarkable preservation of myelin sheaths in the CUP/PLX mice pretreated with PLX. Of additional interest, the diameters of individual axons in the serial block face reconstructions varied quite dramatically along their length, evident in the animations (see supplemental animations) of the reconstructed serial block face images in both control and PLX plus PLX/CUP treated mice. These results indicate a remarkable and previously unappreciated variation in the local diameters of myelinated axons.

### Prophylactic depletion of microglia blocks astrocytosis and OLG loss

Two other key effects of CUP treatment on the CC include oligodendrocyte cell death and astrocytosis (Matsushima G, 2001). Accordingly, we examined whether microgliosis is upstream of these pathological features of CUP or, alternatively if they are independent of microglia and may result directly from OLG damage due to cuprizone. We examined the numbers of OLG, identified with the CC1 antibody, in mice treated with CUP vs. mice pre-treated with PLX and then placed on the combo CUP/PLX diet. Depletion of microglia in the later treatment prevented much of the loss of oligodendrocytes (OLs) in the CC compared to mice on CUP alone (Fig. 5F and G). Similarly, mice fed with CUP alone exhibited a striking increase in GFAP^+^ astrocytes whereas PLX treatment followed by CUP/PLX greatly reduced microgliosis and the associated reactive astrocytosis. A linear regression analysis of microgliosis (Fig. 5D) and astrocytosis (Fig. 5E) vs. demyelination demonstrates strong statistical correlations between these 2 cell types and demyelination in CUP, R^2^ = 0.9691 and 0.9649, respectively. These findings place microglia as essential effectors of the demyelination, OLG death, and the astrocytosis initiated by CUP.

**Figure 5.**
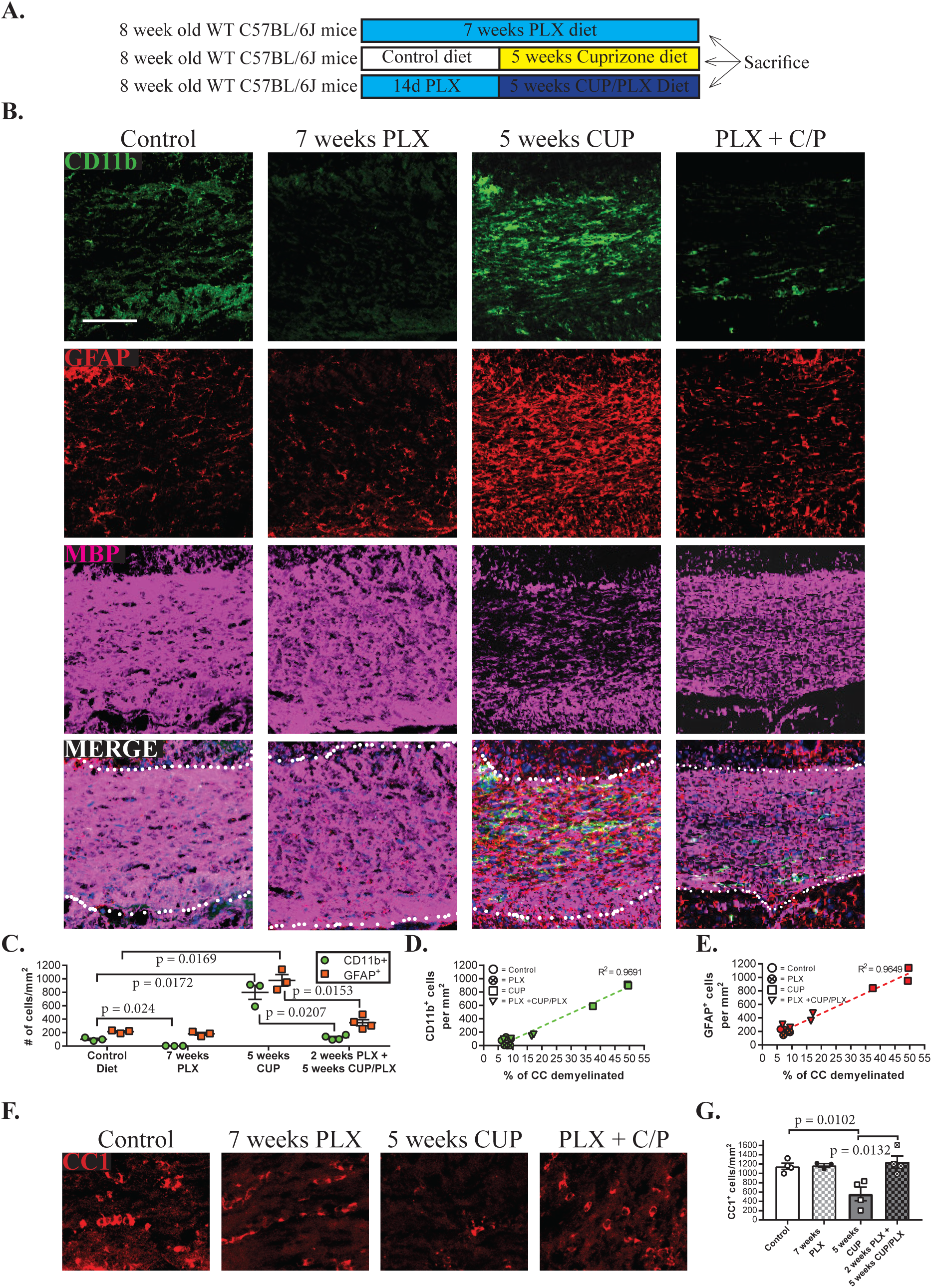
Prophylactic depletion of microglia prevents CUP-mediated astrocytosis and oligodendrocyte loss. A. Schematic of the experimental design, with WT C57BL/6J mice on control diet or treated with PLX for seven weeks, with CUP for 5 weeks, or with PLX for 2 weeks then CUP/PLX for 5 weeks. Mice were then sacrificed and analyzed by immunostaining.
B. Confocal micrographs from each of the cohorts listed above, were stained for CD11b as a marker of microglia, GFAP for reactive astrocytes, and MBP. Scale bar = 200 μm.
C. Quantification of the numbers of microglia (reported by CD11b^+^) and reactive astrocytes (reported by GFAP^+^) per mm^2^ for each of the experimental groups noted including controls (N=3), 7 weeks of PLX (N=3), 5 weeks of CUP (N=3), and 2 weeks PLX + 5 weeks CUP/PLX (N=4). Each symbol represents 1 animal; data are mean +/-s.e.m.
D. Linear regression correlating the numbers of CD11b^+^ cells to the extent of demyelination based on the quantification shown in panels 5B and C. R^2^ = 0.9691
E. Linear regression correlating the numbers of GFAP^+^ cells to the extent of demyelination in the CC based on the quantification shown in panels 5B and C. R^2^ = 0.09649
F. Microglia are required for the loss of oligodendrocytes that results from cuprizone treatment. Confocal micrographs of the CC stained for the mature oligodendrocyte marker CC1 from control mice and from mice treated with PLX (7 weeks), CUP (5 weeks), and PLX + CUP/PLX (2 + 5 weeks). Oligodendrocytes are lost in CUP treated but not PLX + CUP/PLX treated mice.
G. Quantification of the numbers of oligodendrocytes (CC1+ cells) per mm^2^ from the experimental groups shown in Panel F. Each symbol represents 1 animal; data are mean +/- s.e.m.

### Microglia activation is sufficient to induce demyelination

Our results indicate that microglia contribute to demyelination in the CUP model. They are consistent with a two-hit model in which damaged myelin/OLGs require a second microglial-dependent mechanism to undergo removal. They raise the possibility however that activation of microglia by itself (without CUP-mediated damage) may be sufficient to drive demyelination. To address this possibility, we examined the effect of focally activating microglia in the healthy CNS using CSF1, previously used in a pain model (Guan et al., 2015). To this end, we stereotactically injected 1 μl (30 ng/μl) of recombinant colony stimulating factor 1 (rCSF1) into the CC of both *CX3CR1*^*CreER-iresGFP/*^*;Rosa26*^*stop-DsRed*^ (Fig. 6A) and WT (Fig. 6C) mice on either control or a PLX diet. Injection of rCSF1 in the CC resulted in focal microgliosis and associated demyelination, as evident by staining for GFP, DsRed, and MBP, respectively (Fig. 6B). In WT mice, CD11b immunoreactivity was used to determine microgliosis (Fig. 6D and E). In contrast, no microgliosis or demyelination was detected in mice pretreated and maintained on PLX at the time of CSF injection (Fig. 6B-F), indicating demyelination required microglia activation and was not an effect of injecting CSF1 on myelin per se. This also agrees with data indicating that the CSF1 receptor, i.e. *CSF1r*, is expressed by microglia but not by other cells in the adult CNS (Chitu & Stanley, 2017; Y. Zhang et al., 2014). We confirmed these CD11b cells were indeed microglia, and not infiltrating macrophages, by fate mapping cells in *CX3CR1*^*CreER-iresGFP/+*^*;Rosa26*^*stop-DsRed*^ mice (Fig. 6B). Thus, in both vehicle injected and CSF1 injected mice, the CC only contained predominantly GFP^+^/DsRed^+^ cells, identifying them as resident microglia (Fig. 4B). The rCSF1 injected mice on PLX diet did contain a few GFP^+^/DsRed^-^ macrophages with no demyelination (Fig. 6B), likely associated with an injury response at the injection site. Together, these data provide strong support that direct activation of resident CNS microglia is sufficient to induce demyelination.

**Figure 6.**
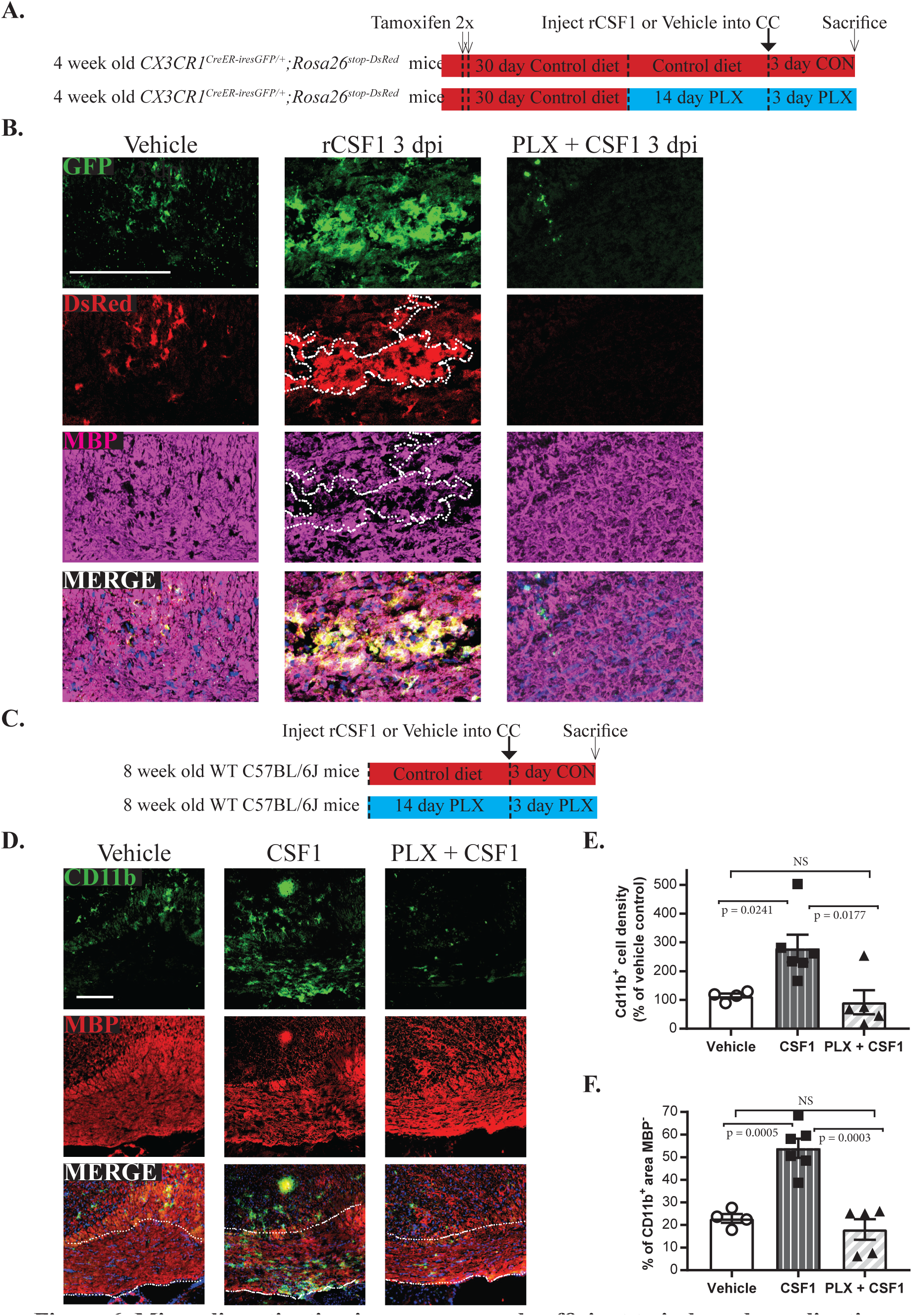
Microglia activation is necessary and sufficient for demyelination. A. Schematic of the experimental design for the stereotactic injection of vehicle or recombinant CSF1 into the CC of *CX3CR1*^*CreER-iresGFP/*^*;Rosa26*^*stop-DsRed*^ mice on a control diet or of CSF1 into the CC of mice pretreated with PLX (2 weeks) and then maintained on PLX for 3 days.
B. Corresponding confocal micrographs from the CC of the experimental groups described in panel A stained for GFP, DsRed, and MBP. Injection of rCSF1 elicits a focal microgliosis evidenced by DsRed+ cells in mice on a control but not PLX diet. There is a corresponding area of focal demyelination associated with the microgliosis induced by CSF1. Scale bar =100 μm.
C. Schematic of the experimental design for injection of CSF1 into the CC of WT mice on control or pretreated with PLX (2 weeks) and then maintained on PLX for 3 days.
D. Confocal micrographs from the CC stained for CD11b as a marker of microglia and MBP. Scale bar = 100 μm.
E. Quantification of the numbers of CD11b^+^ cells in focal lesions resulting from injections of vehicle (N=4) or rCSF1 (N=6) into the CC of mice on a control diet or of rCSF1 into the CC of mice on a PLX diet (N=5). Each symbol represents 1 animal. Data are mean +/- s.e.m.
F. Quantification of demyelination, based on MBP levels, induced by CSF1 induced microgliosis in the experimental cohorts described in panel E.

### CSF1 downregulates microglia Tmem119 similar to other models of demyelination

In MS, microglia lose their expression of homeostatic markers (Zrzavy et al., 2017) including transmembrane protein 119 (Tmem119) which is expressed in both mouse and human microglia (M. L. Bennett et al., 2016). Using a rabbit anti-Tmem119 polyclonal antiserum that stains microglia, we next asked if microglia change their expression of Tmem119 in various murine models of demyelination (CUP, lysophosphatidylcholine (LPC)-induced demyelination, and CSF1-induced demyelination). To this end we stained sections for CD11b and Tmem119 from controls, 5 days post LPC injection, and 3 days post CSF1 injection (Fig 7A). Controls exhibited robust co-expression of CD11b and Tmem119 whereas microglia (CD11b^+^ cell) found in lesions sites lost their expression of Tmem119 in all 3 demyelination models (Fig 7A). Additionally, microglia outside of lesions sites in all 3 demyelination models still retained their expression of Tmem119. These results raise the possibility that the phenotype of CSF1 activated microglia may resemble that in other demyelination models and in human MS. Future studies to further delineate their phenotypes will be of great interest in this regard.

**Figure 7.**
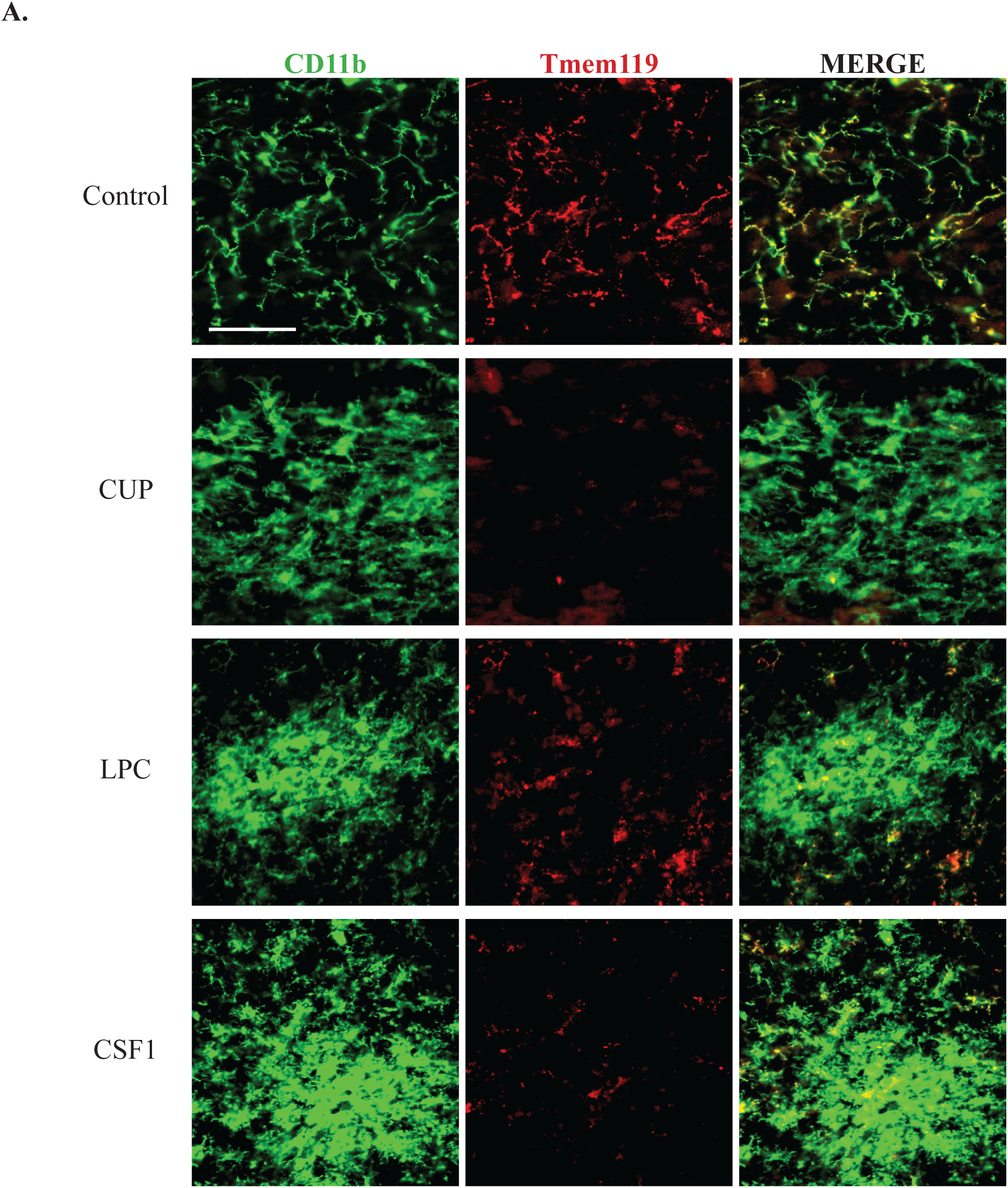
Microglia downregulate Tmem119 in multiple models of demyelination, including CSF1-induced microgliosis. A. Representative confocal micrographs from the CC of mice on control diet, 5 weeks of CUP, 5 days post LPC injection, and 3 days post CSF1 injection. Sections are stained for CD11b and Tmem119. Scale bar = 100 μm.

## DISCUSSION

There is general agreement that microglia contribute to the pathology of both inflammatory demyelination, i.e. EAE (Goldmann et al., 2013; Nissen, Thompson, West, & Tsirka, 2018) and toxin induced demyelination, i.e. cuprizone (Beckmann et al., 2018; Y. Kondo, Adams, Vanier, & Duncan, 2011; Skripuletz, et al., 2010). Their precise role and significance in demyelination has been unclear. In addition, it has been difficult to differentiate the contribution of microglia and infiltrating macrophages in these models as methods to reliably distinguish these cell types have only recently become available. Here, we demonstrate by using a fate-mapping strategy and pharmacological manipulations that microglia are necessary for CUP-mediated demyelination and that their activation is sufficient to drive demyelination. These results, which highlight a key role of microglia in demyelination, are considered further below.

### Microglia but not infiltrating macrophages are found in CUP lesions

Using differential fate mapping (Fig. 1), we show that microglia, not infiltrating macrophages, are found in CUP lesions; if there is a peripheral contribution in the CUP model, it is minimal. In further support of these results, there were no differences in the numbers of CD11b+ cells in the CC or the extent of demyelination between WT and CCR2-/- mice on CUP (Supplemental Fig. 3). These results differ from previous studies that reported modest (McMahon EJ, 2002) or significant (Lampron et al., 2015) macrophages in the CUP model; in the later case, peripheral macrophage infiltration was CCR2-dependent but did not impact the extent of demyelination. These studies used whole body irradiation followed by reconstitution of the bone marrow from a GFP+ donor to address the contribution of macrophages. Whole body irradiation is known to disrupt the blood brain barrier and increase permeability in rodents and humans (Mildner et al., 2007; Nakata H, 1995; Qin, Zheng, Tang, Li, & Hu, 1990), which likely accounts for the macrophage infiltration observed in these studies. Also consistent with disruption of the blood brain barrier is that T-cells were reported in the CC of CUP treated mice following whole body irradiation and bone marrow reconstitution (Remington, Babcock, Zehntner, & Owens, 2007). In contrast to the CUP model, in which the BBB remains intact (A. Kondo, Nakano, & Suzuki, 1987; McMahon EJ, 2002), peripheral blood monocytes do significantly infiltrate and contribute to the demyelinating pathology in other murine models including in EAE (Ajami, et al., 2011; Bourrie et al., 1999) and following stereotactic injection of LPC (de Paula Faria et al., 2014).

### Microglia are necessary for demyelination in the cuprizone model

The cuprizone mouse model is a powerful tool for the investigation of demyelination and has been used for over 50 years (Carlton, 1966, 1967; Suzuki & Kikkawa, 1969). CUP leads to oligodendrocyte specific metabolic dysregulation and selective oligodendrocyte apoptosis, although there is no general consensus of how CUP disrupts oligodendrocytes (Ács & Komoly, 2012; Heather A. Arnett, et al., 2002; Goldberg et al., 2013). Changes in both oligodendrocyte mitochondrial/energy and lipid metabolism have been reported in this model (Praet, et al., 2014).

CUP has been considered to be a direct oligodendrocyte toxin that in turn activates microgliosis. Older studies suggested that microglia may also contribute to CUP-mediated demyelination based on modest beneficial effects of minocycline treatment on disease progression (Pasquini et al., 2007; Skripuletz, et al., 2010). We show here that resident microglia not only contribute but are essential for CUP-mediated demyelination. Coupling PLX with CUP slowed progression of demyelination at three weeks (Fig. 3). Prophylactically depleting microglia further reduced microglia numbers and eliminated most of the rebound microgliosis seen at 5 weeks on the combination diet. Importantly, this prophylactic depletion blocked most of the CUP-mediated demyelination (Fig. 4) demonstrating an unequivocal requirement for microglia in driving CUP-dependent demyelination. These results agree with recent reports in which other CSF1 inhibitors used to deplete microglia resulted in a delay of demyelination mediated by cuprizone (Beckmann, et al., 2018; Wies Mancini, Pasquini, Correale, & Pasquini, 2018) and EAE (Nissen, et al., 2018) further demonstrating the significance of microglia activation as crucial contributors to disease progression.

Given that CUP is thought to target oligodendrocytes specifically, these results support a “two-hit” model in which CUP drives oligodendrocyte and myelin pathology rendering them sensitive to secondary microglia-mediated demyelination and removal of damaged myelin/oligodendrocytes. In support of this model, treatment with cuprizone for just two weeks, which is not associated with evident myelin pathology, can result in subsequent microgliosis and demyelination after another 3 weeks (Doan et al., 2013). In a recent report, a similar brief two week CUP treatment was found to result in myelin hypercitrullination that predisposed the myelin sheath to demyelination when the immune system was activated with adjuvant (Caprariello et al., 2018). We have found that the myelin sheaths that persist after cuprizone treatment in the absence of microglia appear subtly aberrant in their EM appearance (Fig. 4). In initial studies, we have also found that such persistent myelin in treated animals undergoes substantial demyelination with an associated microgliosis, ten days after mice are removed from CUP/PLX and returned to a normal diet. Taken together, the studies indicate that myelin damaged by cuprizone is marked for subsequent destruction and removal by microglia. The signal(s) that mark these damaged myelin sheaths for clearance remain to be identified but are of considerable interest given their potential pathological significance.

Similarly, microglia in the CNS and macrophages in the PNS, also contribute to the pathology of genetically aberrant myelin. Mice deficient for *2’,3’-Cyclic Nucleotide 3’ Phosphodiesterase* (*Cnp*) have minor myelin abnormalities but exhibit low grade inflammation and late stage catatonia (Hagemeyer et al., 2012). Treatment with PLX reduced symptom severity (Janova et al., 2018) implicating microglia in the pathology of genetically sensitive oligodendrocytes and myelin. Additional support for this model is found in Charcot-Marie-Tooth type 1 neuropathy, a genetic mutation specific for myelinating Schwann cells, which is characterized by inflammation, peripheral demyelination, muscle denervation, increased macrophage numbers, and increased expression of CSF1 (Groh et al., 2012), Treatment with PLX reduced macrophage numbers by ∼70% and promoted axonal integrity and muscle innervation (Klein et al., 2015).

Unexpectedly, microglia numbers rebounded in the CC of mice being maintained on the combination CUP/PLX diet. While microglia were substantially depleted at 3 weeks of this diet, two weeks later microglia numbers and the extent of demyelination had substantially increased (Fig. 3). The robust depletion of microglia in the cortex of these mice underscores the efficacy of the PLX treatment in other sites and suggests that microglia in the demyelinating CC specifically become insensitive to the CSF1R inhibitor. Potentially, injury signals that drive microgliosis during demyelination may compensate for CSF1R inhibition and drive its repopulation via rapid proliferation (Monica R. P. Elmore, et al., 2014; Huang, et al., 2018). One such candidate is the lipid sensor Triggering receptor expressed on myeloid cells 2 (TREM2), which is upregulated by microglia in the CC during CUP treatment (Cantoni et al., 2015). TREM2 is activated by phospholipids, such as those from degraded myelin, and can in turn activate the adaptor protein DAP12, promoting microglia survival and expansion (Yeh, Hansen, & Sheng, 2017). TREM2 activation of DAP12 may compensate for the inhibition of CSF1R signaling by PLX which would normally be expected to activate DAP12 (Ulland, Wang, & Colonna, 2015). Future studies will be needed to identify the precise mechanism(s) that drive CC microgliosis during cuprizone treatment.

### Microglia activation is sufficient to induce demyelination

In gain-of-function studies, we further demonstrated that direct activation of microglia by CSF1 injection is sufficient to initiate robust, focal CNS demyelination (Fig. 4). Cells in the lesion were unambiguously identified as microglia by fate mapping and were shown to be essential for demyelination as rCSF1 did not elicit demyelination in microglia depleted mice. Additional evidence for the key role of microglia, and of the CSF1-CSF1R axis, in demyelination, is underscored by mutations in the CSF1R which result in adult onset, hereditary diffuse leukoencephalopathy (Adams, Kirk, & Auer, 2018; Saitoh et al., 2013). These results indicate that microglia activation, either as a primary event or as a consequence of myelinating cell injury, elicits demyelination. They further that suggest microglial activation may likewise directly contribute to demyelination in many myelin disorders.

In addition to alterations in myelination, chronic CUP treatment has been used to model other neurological disorders such schizophrenia and epilepsy (Praet, et al., 2014). Mice on CUP diet exhibit a loss of mossy neurons in the hippocampus, impaired spatial memory, decreased social behavior, and neurotransmitter dysregulation in the prefrontal cortex (Hoffmann, Lindner, Gröticke, Stangel, & Löscher, 2008; Makinodan et al., 2009; Xu, Yang, Rose, & Li, 2011; Xu et al., 2009). Rather than being secondary to the effects of CUP on myelination, our results raise the possibility that microglia activation may also underlie these various pathologies. This is consistent with an emerging role of neuroinflammation in neurobehavioral disorders including obsessive-compulsive disorder, major depressive disorder, schizophrenia, and stress (Calcia et al., 2016; Krabbe et al., 2017; Mondelli, Vernon, Turkheimer, Dazzan, & Pariante, 2017; Setiawan, Wilson, Mizrahi, & et al., 2015).

### Mechanism of microglia-mediated demyelination

Key related questions are how activated microglia recognize damaged myelin and drive demyelination. Activated microglia release inflammatory cytokines that may contribute to oligodendrocyte death and demyelination, consistent with earlier *in vitro* studies (Pasquini, et al., 2007). In addition, microglia have an essential role in clearance of apoptotic cells in the adult CNS (Fourgeaud, et al., 2016; Mazaheri et al., 2014; Sierra, et al., 2010). In some circumstance macrophages/microglia may induce death of damaged but still viable cells - a process termed phagoptosis (Neher, Neniskyte, Hornik, & Brown, 2014; Neher et al., 2011). These microglial mechanisms may thus contribute to the death of oligodendrocytes in the CC that are viable but significantly dysregulated due CUP treatment (Caprariello, et al., 2018) whereas in the absence of microglia, such oligodendrocytes may survive (Fig. 5). In the future, an assessment of the numbers of apoptotic oligodendrocytes with and without microglia (i.e. PLX treatment) will be useful to corroborate this possibility.

In addition to direct toxic effects, microglia are known to activate astrocytes which are candidates to exacerbate the demyelination. In agreement with previous reports (Matsushima G, 2001), CUP treatment resulted in reactive astrocytosis indicated by increased GFAP staining, that is highly correlated to the extent of demyelination and microgliosis (Fig. 5). Microglia were recently shown to directly activate astrocytes via a series of soluble factors, i.e. interleukin1α, tumor necrosis factor and complement component 1, subcomponent q (Liddelow et al., 2017). In agreement, we found blocking microgliosis in the CUP model by PLX proportionately reduced the levels of reactive astrogliosis (Fig. 5). As PLX directly targets microglia via inhibition of CSF1R, a growth factor receptor that is not expressed by astrocytes (Teh et al., 2017), our results indicate astrocytosis in the CUP model is secondary to microglial activation. Other studies strongly implicate astrocytes as crucial amplifiers of inflammation in demyelination (Moreno et al., 2014; Raasch et al., 2011; van Loo et al., 2006), neurodegeneration (Hsiao, Chen, Chen, Tu, & Chern, 2013), and as a source of soluble factors that are toxic for neurons and oligodendrocytes (Liddelow, et al., 2017). Thus, PLX may limit demyelination by reducing both primary (i.e. microglial) and secondary (e.g. astrocytic) sources of oligodendrocyte toxic factors.

The phenotype of microglia is highly plastic (Ransohoff & El Khoury, 2015). Microglia alter their gene expression in development and in models of demyelination and neurodegeneration (Bedard, Tremblay, Chernomoretz, & Vallieres, 2007; Gao et al., 2017; Hammond et al., 2018; Hirbec, Noristani, & Perrin, 2017; Li et al., 2018). The precise effects of rCSF1 onto CC microglia remain an open question. Our own analysis of the phenotype of microglia following CSF1 injection was confined to demonstrating their lack of Tmem119 immunoreactivity. Loss of microglia Tmem119 is shared between all models of demyelination used in this study (CUP and LPC). Brains from patients with Alzheimer’s disease, Parkinson’s disease, and amyotrophic lateral sclerosis contain Tmem119 positive microglia, whereas microglia in active MS lesions lack Tmem119 (Satoh et al., 2016; Zrzavy, et al., 2017).

Of note, increased CSF1 levels have been reported to correlate strongly with microglial activation and neuronal loss in EAE (Gushchina et al., 2018). These latter findings suggest CSF1 may promote EAE pathology either by driving increased microglia numbers and/or their transition to a demyelinating phenotype. Our current studies underscore the striking ability of CSF1 to activate microglia as a primary event in demyelination, independent of oligodendrocyte/myelin damage. Future efforts to further delineate the phenotype of CSF1 stimulated microglia in the CC and how they compare to microglia activated in inflammatory demyelination in EAE and MS will therefore be of great interest

## MATERIALS AND METHODS

### Animals and administration of tamoxifen

All rodent experiments were carried out under an approved animal protocol in accordance with the guidelines of the Institutional Animal Care and Use Committee of New York University School of Medicine. Mice were maintained under standard husbandry conditions at an NYU vivarium. Details on all strains used can be found in Table 1.

**Table 1.**
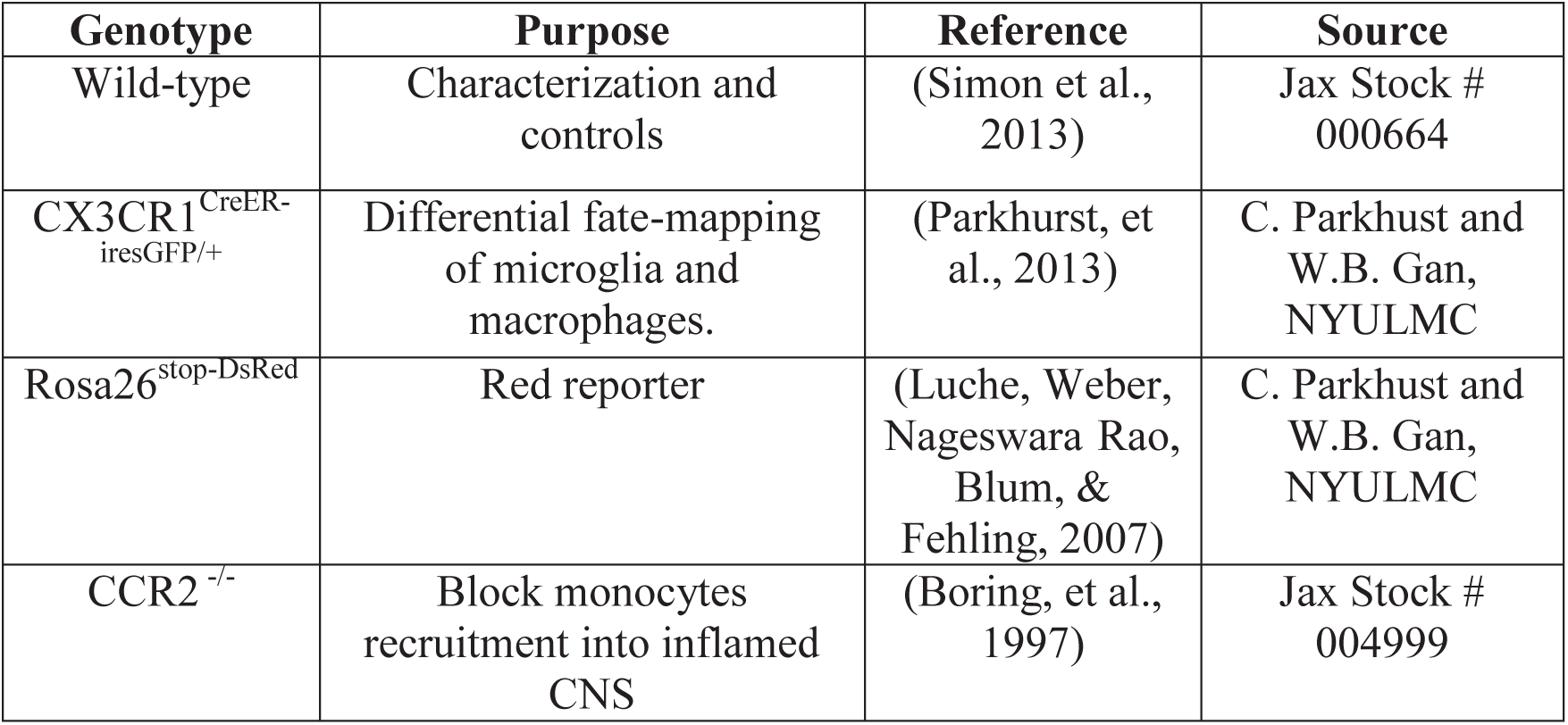
Summary of Mouse Strains

### Genotyping

Small (∼0.5 cm) tail trimmings from mouse pups (<21 day old) were collected and placed in eppendorf tube. 200 μl of tail lysis buffer (500ml distilled water, 18ml of 1.5M Tris-HCl (pH 8.8), 10.8 ml of 5 M NaCL, 1.08 ml of 0.5 M EDTA, and 13.5 ml of Tween-20 (Sigma) was added to each tube along with 4μl of Proteinase K (Sigma). Tails were incubated overnight at 55° C. Tubes were then placed at 100° C for 15 minutes to heat inactivate Proteinase K. 1-2μl of solution was used for each PCR reaction. PCR reaction solution which included: 9μl of nuclease free water, 12μl of Dreamtaq polymerase (Thermofisher), and 1μl of both forward and reverse primers. Samples where then placed into Arktik Thermal Cycler (Thermofisher). Samples were then run on 2.0% agarose gel. Details of PCR primer sequences, thermocycler protocols, and expected band sizes can be found in Table 2. For all reactions a no DNA negative control and WT DNA control sample were run in tandem.

**Table 2.**
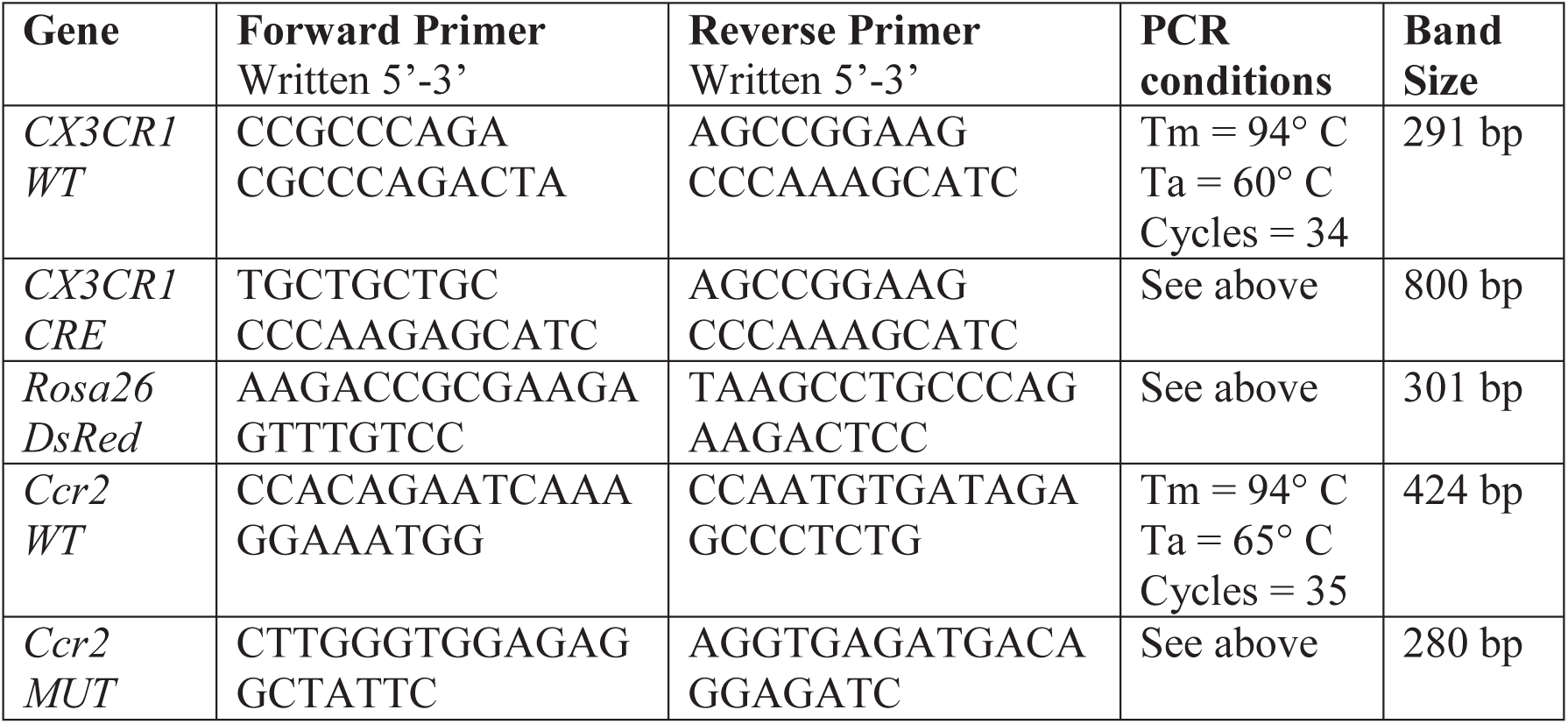
Genotyping information

Differential labeling of microglia and peripheral macrophages using tamoxifen

*CX3CR1*^*CreER-iresGFP/+*^ mice were crossed with *Rosa26*^*stop-DsRed*^ line. Four to six week old *CX3CR1*^*CreER-iresGFP/+*^*;Rosa26*^*stop-DsRed*^ mice were given 10 mg tamoxifen (Sigma) in corn oil by oral gavage on alternate days for a total of 2 gavages. Mice were given 30 days following the final administration of tamoxifen to allow for the loss of DsRed labeling in peripheral immune cells before initiating experimental procedures (Parkhurst, et al., 2013).

### Cuprizone model of demyelination

Adult (8-10 week old) mice were fed 0.2% cuprizone (sigma) in irradiated feed prepared by Research Diets. 3 and 5 weeks of cuprizone diet were taken as approximate disease onset and disease peak, respectively (Wergeland, et al., 2012).

### Colony stimulating factor 1 experiments

1 μl of recombinant CSF1 (Gibco), diluted to 30 ng/ml in 0.1% BSA, was stereotactically injected in the corpus callosum using coordinates 0.79 mm lateral and1.86 mm ventral to the bregma. Cortical injections were done 0.79 mm lateral and 0.89 mm ventral to the bregma.

### Pharmacological depletion of microglia

The CSF1 inhibitor PLX3397 (Plexxikon) was mixed into AIN-76A standard irradiated chow by Research Diets at a dose of 290mg/kg of chow (Monica R. P. Elmore, et al., 2014). Mice fed this diet underwent targeted and reversible depletion of microglia. Research Diets also combined PLX3397 with 0.2% cuprizone in standard irradiated chow to make PLX/CUP chow.

### Tissue processing

Mice were deeply anesthetized with an intraperitoneal injection of 0.1 ml of Beuthanasia (Merck). Upon loss of paw and tail reflex, mice were mounted to dissection board and underwent transcardial perfusion with 30 ml of cold PBS followed by 30 ml of cold 4% PFA (Electron Microscopy Sciences). Brains were removed and post fixed for four hours in 4% PFA then equilibrated with to 30% sucrose (Fisher Scientific) overnight at 4° C. Brains were then embedded in O.C.T. Compound (Scigen) and placed at −80° C overnight before processing. Once brains were completely frozen, they were placed at −20° C and mounted on a Leica CM3050. 20 μm thick coronal sections were taken through the corpus and hippocampus. Sections were mounted onto Colorfrost Plus microscope slides (Fisher Scientific) and left to dry overnight before being stored at −80° C.

### Immunofluorescence

Slides were washed three times with PBS for 5 minutes before being blocked for 1 hour at room temperature in blocking solution (10% normal goat serum, 1% BSA, 0.25% Triton X-100 in PBS). Slides were then incubated with primary antibodies overnight at 4° C in antibody solution (1% BSA, 0.25% Triton X-100 in PBS). Slides underwent three washes with PBS at room temperature. Secondary labeling with Alexa Fluor antibodies (Invitrogen; 488, 594, 680; 1:1000 dilution) raised in goat was carried out for 1 hour at room temperature. Slides were then washed three times in PBS and incubated in Hoechst 33258 (Invitrogen) for 5 minutes diluted 1:5000 in PBS. Slides were washed three times in PBS and three more times in distilled water, before being cover-slipped using Fluoroshield (Sigma). Slides were dried overnight in the dark prior to imaging.

Primary antibodies used in these experiments included: rat anti-CD11b (BD Biosciences) 1:400, mouse anti-GFAP (Sigma), 1:400, rabbit anti-GFAP (Dako), 1:400, rabbit anti-GFP (Invitrogen), 1:1000, chicken anti-GFP (Invitrogen), 1:750, rabbit anti-RFP (Rockland Immunohistochemicals), 1:750 which recognizes DsRed, mouse anti-MBP (Millipore) 1:500, and rabbit anti-TMEM119 (gift from M. Bennett and B. Barres, Stanford)

### Fluorescent imaging

Fluorescent images were obtained as Z-stacks of 1 μm optical sections using a confocal laser-scanning microscope (LSM 800, Zeiss). For experiments analyzing demyelination, sections spanning the entire CC were surveyed for demyelinated lesions. Then using the 10x objective, five images per mouse were taken of CC of demyelinated regions, taken as the largest regions with diminished MBP fluorescence. A similar strategy was used to determine cell numbers using the 20x objective to aid in quantification. Imaging was performed in a blinded manner.

### Quantitative analyses of cell numbers and myelin

For analyses of CC myelin, images were minimally processed for brightness and contrast in Fiji (Schindelin et al., 2012), with all samples in an experiment adjusted equivalently. The CC was then traced, and the percentage of total area of MBP was measured. The cell counter tool was used to determine cell numbers. For all studies, five sections per mouse were quantified in a blinded manner. Data from three to five mice were combined to determine the average and standard error of means (SEM) using GraphPad Prism version 7.00. Data are plotted as averages ± SEM. Student’s t-test was performed to calculate P values.

### Electron Microscopy

Male C57BL/6 mice (control and treated with PLX for two weeks and then PLX/CUP) were subjected to trans-cardiac perfusion with 4% PFA, 2.5% glutaraldehyde, and 0.1M sucrose in 0.1M phosphate buffer (PB, pH 7.4). The body of corpus callosum was later excised from the dissected brain, and the tissue was fixed in the same fixative solution, followed by a PB containing 2% OsO4 and 1.5% potassium ferrocyanide for 1 hour. The tissues were then stained with 1% thiocarbohydrazide (Electron Microscopy Sciences, EMS, PA) for 20 min, 2% osmium tetroxide for 30 min, and 1% aqueous uranyl acetate at 4° C overnight. An En Bloc lead staining was performed at 60° C for 30 min to enhance membrane contrast. Brain samples were dehydrated in alcohol and acetone, and embedded in Durcupan ACM resin (EMS, PA) (Wilke et al., 2013). Tissue samples were analyzed with a scanning electron microscope (SEM) (Zeiss Gemini 300 SEM with 3View). Cropped regions consisting of 100 slices each 100 nm apart, that were 16 μm × 16 μm, with a pixel size of 4 nm providing a voxel resolution of 4 nm × 4 nm × 100 nm. Fixed sections from comparable fixed mice were also examined by TEM as previously described (Einheber, Schnapp, Salzer, Cappiello, & Milner, 1996). Tissue sections and imaging were carried out by NYU Langone’s Microscopy Laboratory.

## Supporting information

Serial Block EM recon - Control CC

Serial Block EM recon - 2wkPLX + 5wkC/P CC

## Acknowledgements

We thank C. Parkhurst and W. Gan for advice during the course of this project, M. Bennett for providing antibodies, J. Gallagher, C. Bennett and S. Liddelow for comments on the manuscript, B. Zotter for technical advice, A. Liang, K. Dancel-Manning, C. Petzold, and J. Sall of NYU’s Microscopy laboratory for expert assistance with electron microscopy, and K. Lee for assistance with determining g ratios. This research was supported by grants to J.L.S. from the NYS DOH Stem Cell and the NIH. DM was a recipient of an NRSA fellowship from NINDS.

## Author Contributions

Conceived and designed the experiments: DEM, JLS. Performed the experiments: DEM Analyzed the data: DEM, JLS. Wrote the paper: DEM, JLS. Conceptualized the research: DEM, JLS. Provided project oversight: JLS. Provided the CSF1R inhibitor PLX3397: BLW.

---

**Supplementary Figure 1.**
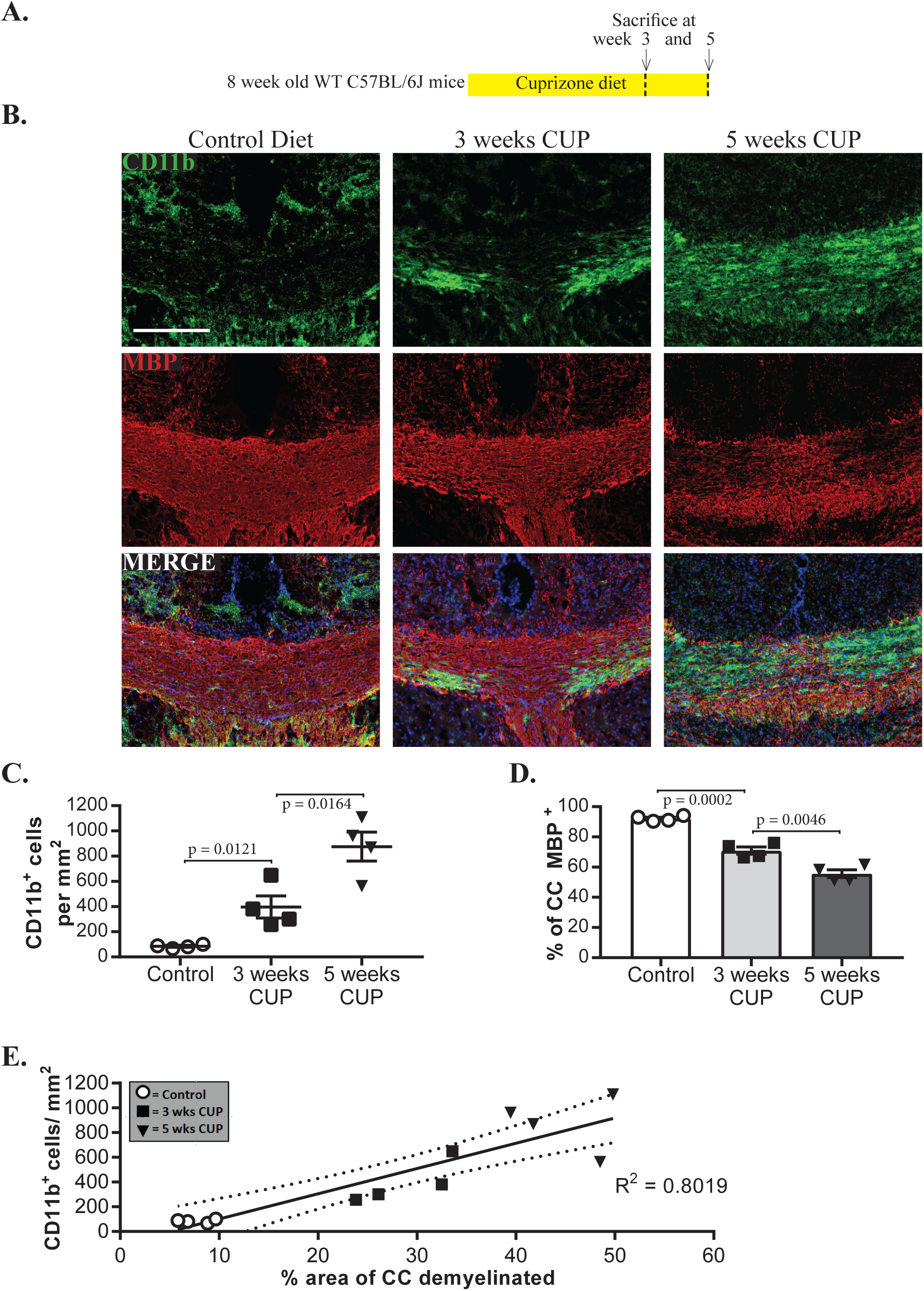
Microgliosis is strongly correlated with demyelination. A. Schematic of the experimental design. Wild type C57BL/6j mice were place on cuprizone diet (CUP) before evaluation of demyelination and microgliosis.
B. Confocal micrographs of coronal sections stained for CD11b and MBP. Merged images also show staining with Hoechst nuclear stain. Scale bar = 200 μm.
C. Quantification of CD11b^+^ cells per mm^2^ shown in B. Each symbol represents 1 animal; N= 4 per group. Data are mean +/-s.e.m.; Student’s t-test.
D. Quantification of percentage of CC positive for MBP; experimental groups correspond to those shown in panel B.
E. Linear regression correlating CD11b^+^ cell density (panel C) to the extent of demyelination (panel D). Control diet (empty circles), 3 weeks CUP (empty squares), and 5 weeks CUP (filled squares) are shown. Dashed lines represent 95% confidence interval. R^2^ value = 0.8019

**Supplementary Figure 2.**
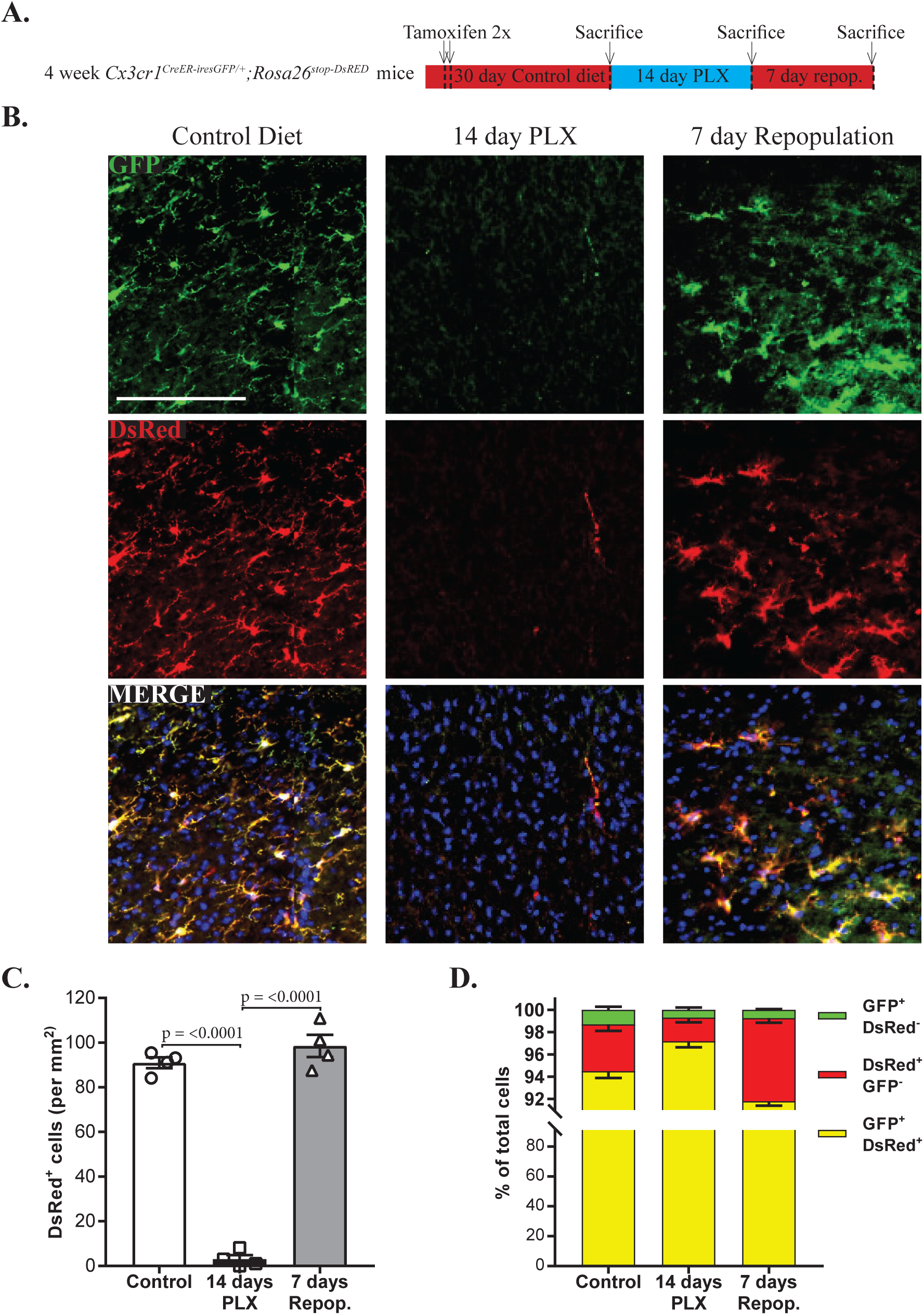
Microglia that repopulate the CNS after PLX3397 depletion are derived from residual, surviving microglia. A. Schematic of the experimental design. *CX3CR1*^*CreER-iresGFP/+*^*;Rosa26*^*stop-DsRed*^ mice were placed on diet containing the CSF1R inhibitor PLX3397 diet (PLX) and evaluated for microglia depletion at two weeks or repopulation after resuming the control diet for an additional week.
B. Confocal micrographs from the cortex stained for GFP and DsRed for each group. Merged images also show staining with Hoechst nuclear stain. Scale bar = 100 μm.
C. Quantification of DsRed^+^ cells/mm^2^ from experimental groups shown in B. Each symbol represents 1 animal; N = 4 per group. Data are mean +/-s.e.m.; Student’s t-test.
D. Quantification of macrophages (GFP^+^ only; green) vs. microglia (DsRed^+^ only, red and GFP^+^ DsRed^+^ double positive; yellow) from experimental groups shown in C.

**Supplementary Figure 3.**
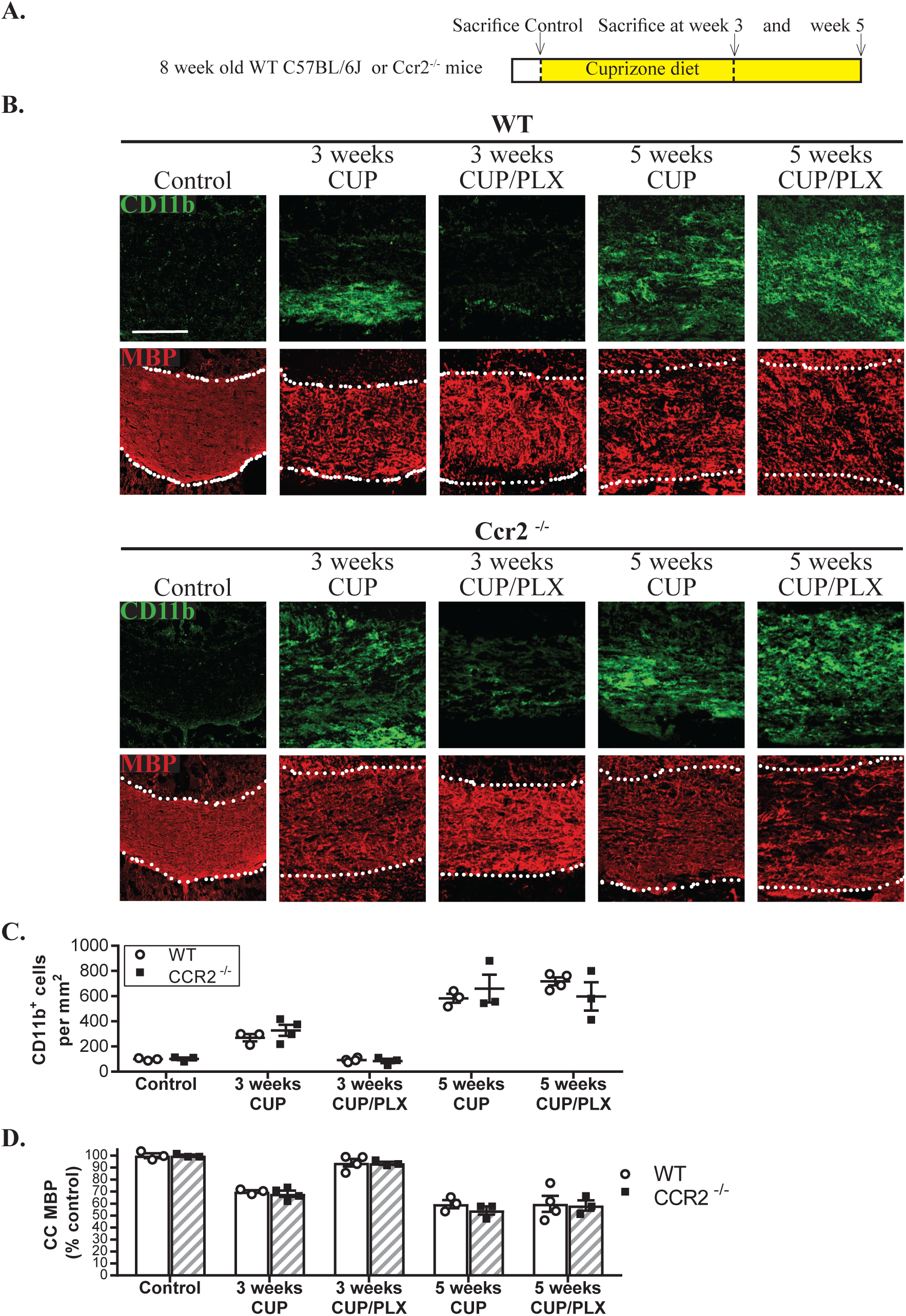
Peripheral macrophages do not contribute to demyelination in the cuprizone model. A. Schematic of the experimental design. 8 week old WT and CCR2 ^-/-^ mice were placed on diets containing CUP for 3 and 5 weeks or CUP/PLX for 3 and 5 weeks before evaluation of the extent of demyelination and microgliosis.
B. Confocal micrographs of the CC from the experimental groups indicated were stained for CD11b and MBP. Scale bar = 100 μm.
C. Quantification of CD11b^+^ cells/mm^2^ from the groups shown in B. WT cohorts included controls (N = 3), 3 weeks CUP (N = 3), 3 weeks CUP/PLX (N = 4), 5 weeks CUP (N = 3) and 5 weeks CUP/PLX (N = 4). For CCR2^-/-^ mice, groups included controls (N = 3), 3 weeks CUP (N = 4), 3 weeks CUP/PLX (N = 3), 5 weeks CUP (N = 3), and 5 weeks CUP/PLX (N = 3). Each symbol represents 1 animal. Data are mean +/-s.e.m.; Student’s t-test. N’s = 3, 4, 3, 3, 3
D. Quantification of CC MBP relative to control diet for the experimental groups of mice shown in B and C.

